# Spatial distribution of benthic macrofauna in the Central Arctic Ocean

**DOI:** 10.1101/353060

**Authors:** Andrey Vedenin, Manuela Gusky, Andrey Gebruk, Antonina Kremenetskaia, Elena Rybakova, Antje Boetius

## Abstract

Permanent ice coverage and the low primary production in the mostly ice-covered Central Arctic ocean basins result in significantly lower biomass and density of macrobenthos in the abyssal plains compared to the continental slopes. However, little is known on bathymetric and regional effects on macrobenthos diversity. This study synthesizes new and available macrobenthos data to provide a baseline for future studies of the effects of Arctic change on macrofauna community composition in the Arctic basins. Samples taken during three expeditions (in 1993, 2012 and 2015) at 37 stations on the slope of the Barents and Laptev Seas and in the abyssal of the Nansen and Amundsen Basins in the depth range from 38 m to 4381 m were used for a quantitative analysis of species composition, abundance and biomass. Benthic communities clustered in five depth ranges across the slope and basin. A parabolic pattern of species diversity change with depth was found, with the diversity maximum for macrofauna at the shelf edge at depths of 100-300 m. This deviates from the typical species richness peak at mid-slope depths of 1500-3000 m in temperate oceans. Due to the limited availability of standardized benthos data, it remains difficult to assess the massive sea ice retreat observed in the past decade has affected benthic community composition. The polychaete *Ymerana pteropoda* and the bryozoan *Nolella* sp. were found for the first time in the deep Nansen and Amundsen Basins, as a potential first sign of increasing productivity and carbon flux with the thinning ice.

## Introduction

The deepest parts of the Central Arctic, including the abyssal plains of the Nansen and Amundsen Basins separated by the Lomonossov Ridge, are among the least studied areas of the global oceans. Most surveys of Arctic benthos have been conducted on the shelf or upper slope, whereas data from the deep basins remain rare and scattered in time and space. First data on the deep-sea fauna of the Central Arctic were obtained in 1935 using trawls and dredges not suitable for taking quantitative samples. Qualitative descriptions of the Central Arctic bathyal and abyssal fauna were made by Soviet expeditions on the vessels *Sadko* in 1935-1937 and *F. Litke* in 1955 [1,2]. First quantitative data using mini-LUBS corer and “Okean” grabs were collected in the 1970s in the Canadian Basin from the American Fletcher’s Ice Island [3] and the Soviet drifting station *North Pole-22* [4]. The authors reported extremely low abundances at depths >1000 m of a few individuals per m^2^ and hence a very low benthos biomass of around 0.04 g/m^2^. A recent review of the Arctic macrobenthos confirms the strong decline in biomass by over an order of magnitude from the outer shelves to the inner basins of the Arctic Ocean [5]. The Arctic deep-sea benthos has likely evolved from shallow-water relatives inhabiting the large shelves, with cold temperatures close to freezing point prevailing across the entire depth range. Till today, the communities of the deep basins and the outer shelf share more than half of their taxa [5], and food limitation seems to be the main factor structuring community composition [6].

Today, when the rapid warming and sea ice decline are likely to change the Arctic ecosystem in its entity, comparative analyses of macrofauna community composition, diversity and abundance in space and time remain challenged by the paucity of quantitative, standardized macrofauna data. Sampling surveys were made during expeditions of the research icebreakers *Ymer*, *Polarstern*, *Oden* and *Polar Sea* in the 1990s and 2000s, but unfortunately not with internationally standardized procedures [7–12]. Over 25 macrobenthic samples were retrieved in 1991 during the *Polarstern* expedition to the Nansen and Amundsen Basins [13,14], but the low volume of sample obtained caused problems in diversity detection [15]. During the *Polarstern* expeditions in 1993 and 1995, nine bathymetric transects were made at the northern slopes of Barents, Kara and Laptev Seas from shelf to abyssal depths. Results of these surveys were partly published by [11], [15] and [16]. The present study provides additional analyses of legacy samples from these expeditions and adds a substantial number of new data from surveys during the sea-ice minimum in 2012 [6,17].

Patterns of bathymetric distribution of benthic fauna were previously reviewed for different temperate and tropical ocean regions [18–23]. According to generally recognised patterns, the density and the biomass of macrobenthos decrease with depth due to the declining flux of particulate matter as main food supply. In contrast, species diversity shows a parabolic pattern with a maximum at depths of 1500-3000 m [24–27]. In the Arctic, the known species diversity of around 1100 taxa does not appear to follow this pattern, and no mid-depth peak of diversity was yet detected, potentially due to the strongly limited food supply (summarised in [28]).

This hypothesis is supported by recent studies of horizontal distribution patterns of macrofauna within the deep-sea Central Arctic. From the shelf to the deep-sea basins, food supply is declining not only because of increasing water depth, but also because northwards, the sun angle and the sea-ice conditions limit light availability to primary producers. A summary of the abundance, biomass and modelled productivity of the Central Arctic macrofauna was recently published by [6], covering samples from a 20-year period and a depth range of 520-5420 m. It was shown that standing stock and production of the benthic communities are several times higher under the seasonal ice than under the multiyear ice zones. This correlation was explained by the different particle flux under the seasonally and permanently ice-covered areas [6,17].

In the present study we aimed at further testing the effect of location, water depth, sea ice cover and phytodetritus flux for a large standardized data set of macrofauna at high taxonomical resolution, focusing on the Eurasian slope and basin. [28] noted before, that analytic differences among investigators make difficult or even impossible to synthesize species lists from different studies. In our study the entire set of samples was identified by the same specialists. Another aim was to test the entire data set available for any indication of decadal change in community composition between the 1990s and today, i.e. before and after the significant sea ice decline in this region [29]. We examined the bathymetric distribution patterns of macrofauna along the Barents and Laptev Sea slopes and the spatial distribution pattern within central parts of the Amundsen and Nansen Basins. Our hypotheses were the following: 1) Variations in macrobenthos distribution are controlled by food availability; 2) Community similarity is high across all depth zones; 3) Sea ice retreat leads to change in the community structure.

## Materials and methods

### Sampling

Material for this study was obtained from three expeditions. During the RV *Polarstern* expedition ARK-IX/4 (August-September 1993), 44 benthic stations were sampled on northern slopes of the Barents and Laptev Seas and adjacent shelf areas and the Nansen Basin. Sampling gears included the multibox-corer (MG) [30] and the 0.25 m^2^ giant box-corer (GKG). From one to fourteen subsamples of the size 0.023 m^2^ were taken from each MG and/or from one to four subsamples of the size 0.022 m^2^ from each GKG using the rectangular frame. Part of these samples was processed previously at the Zoological Museum in St. Petersburg, Russia, and results were published by [15]. The material from additional 27 stations sampled in 1993 has now been analyzed and is used in the present study to obtain a more comprehensive set of baseline data. Information on these stations, including the number of subsamples and sampling area, are shown in Table 1, the entire dataset can be downloaded from doi.pangaea.de/10.1594/PANGAEA.890152. The uppermost 1 cm of sediment from each subsample was washed through the 0.2 mm sieve; the deeper 5 cm were washed through the 0.5 mm sieve. The washed subsamples were preserved in 4% formaldehyde solution. For additional information see [9].

**Table 1.**
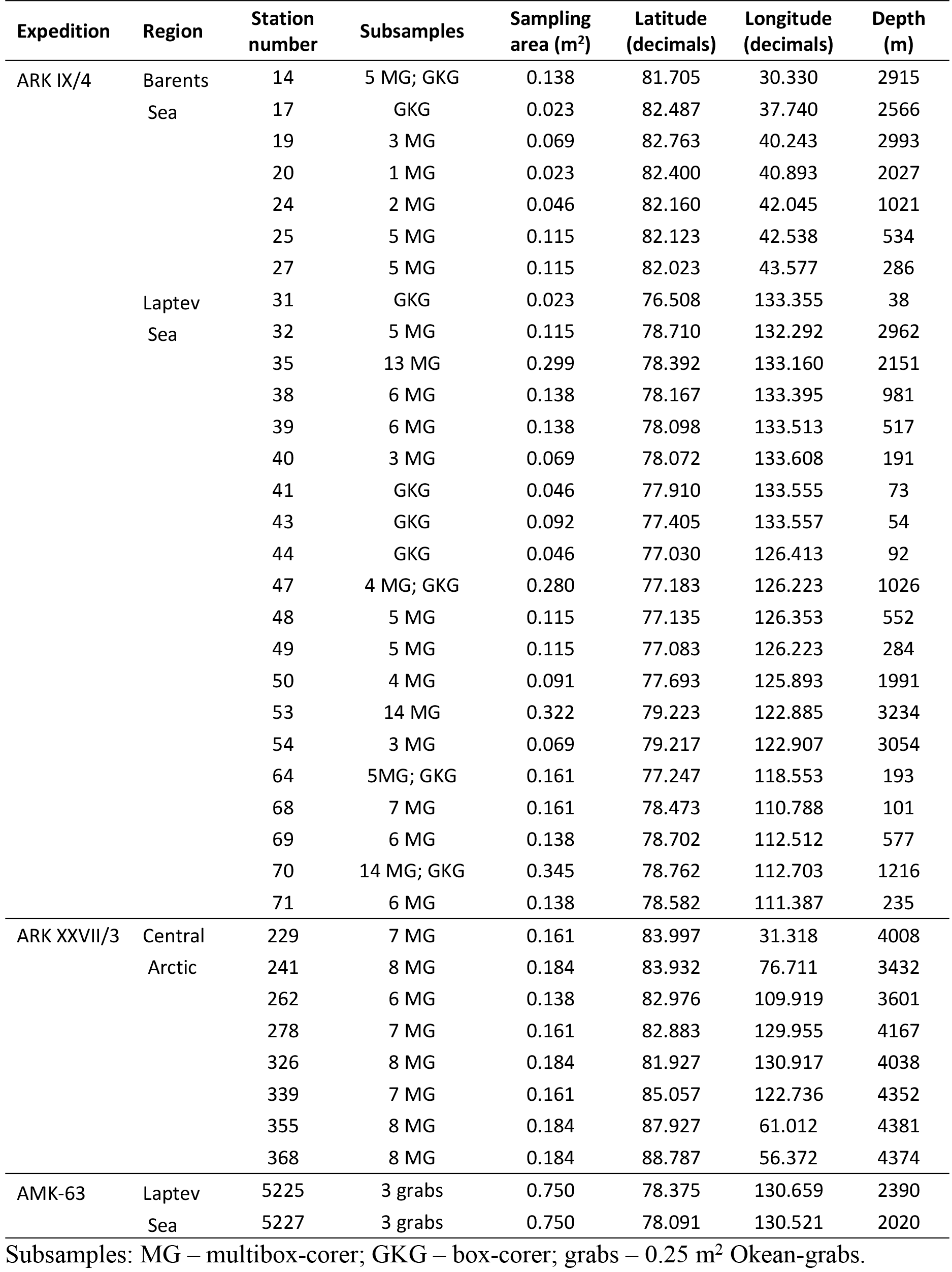
Station data showing the expedition, geographical position, depth, number of subsamples and sampling area at each station.

A total of eight macrobenthic stations were taken in the Nansen and Amundsen Basins during the RV *Polarstern* expedition ARK-XXVII/3 (August-September 2012). Sampling of macrofauna was conducted by the multibox-corer with six to eight 0.023 m^2^ subsamples taken at each station [30]. Subsamples were washed separately through the 0.5 mm mesh size sieve. Four cores were fixed with 4% formaldehyde and two to four cores (depending on the number of successfully closed cores) were fixed with 96% ethanol. Data on the biomass and density of macrobenthos based on this material were published by [6]. Our current study adds the taxonomic analysis. For additional information on sampling procedures see [31].

Samples taken during the Cruise 63 of *Akademik Mstislav Keldysh* (September 2015) at the base of the Laptev Sea slope were also examined. Two stations using an *Okean* grab sampler with 0.25 m^2^ sampling area [32] were performed with three grab replicates per station. Each grab sample was washed through the 0.5 mm sieve and fixed by 5% formalin.

For all samples retrieved in 1993 and 2012 we assessed factors governing variations in macrobenthos composition including porosity, chloroplast pigments concentrations, protein and phospholipids contents, bacterial abundance and activity (esterase and lipase activity) and organic carbon (doi.pangaea.de/10.1594/PANGAEA.890152). For methods see [33], and [34]. Samples for these parameters were obtained by multi-corer samples to retrieve the undisturbed top centimeter layer of sediment at the box core stations during the same expeditions. In addition we used the sea-ice coverage data measured at the moment of sampling at each station. Additional data from the Ocean Floor Observation System (OFOS) used in the ARK-XXVII/3 expedition allowed estimating the area of the *Melosira arctica* algal patches on the sediment surface (https://doi.pangaea.de/10.1594/PANGAEA.803293; For methods see [9] and [31].

The map of the sampling area with stations is shown in Fig 1. Station data are given in Table 1. The environmental data used in this study can be downloaded from doi.pangaea.de/10.1594/PANGAEA.890152.

**Fig 1.**
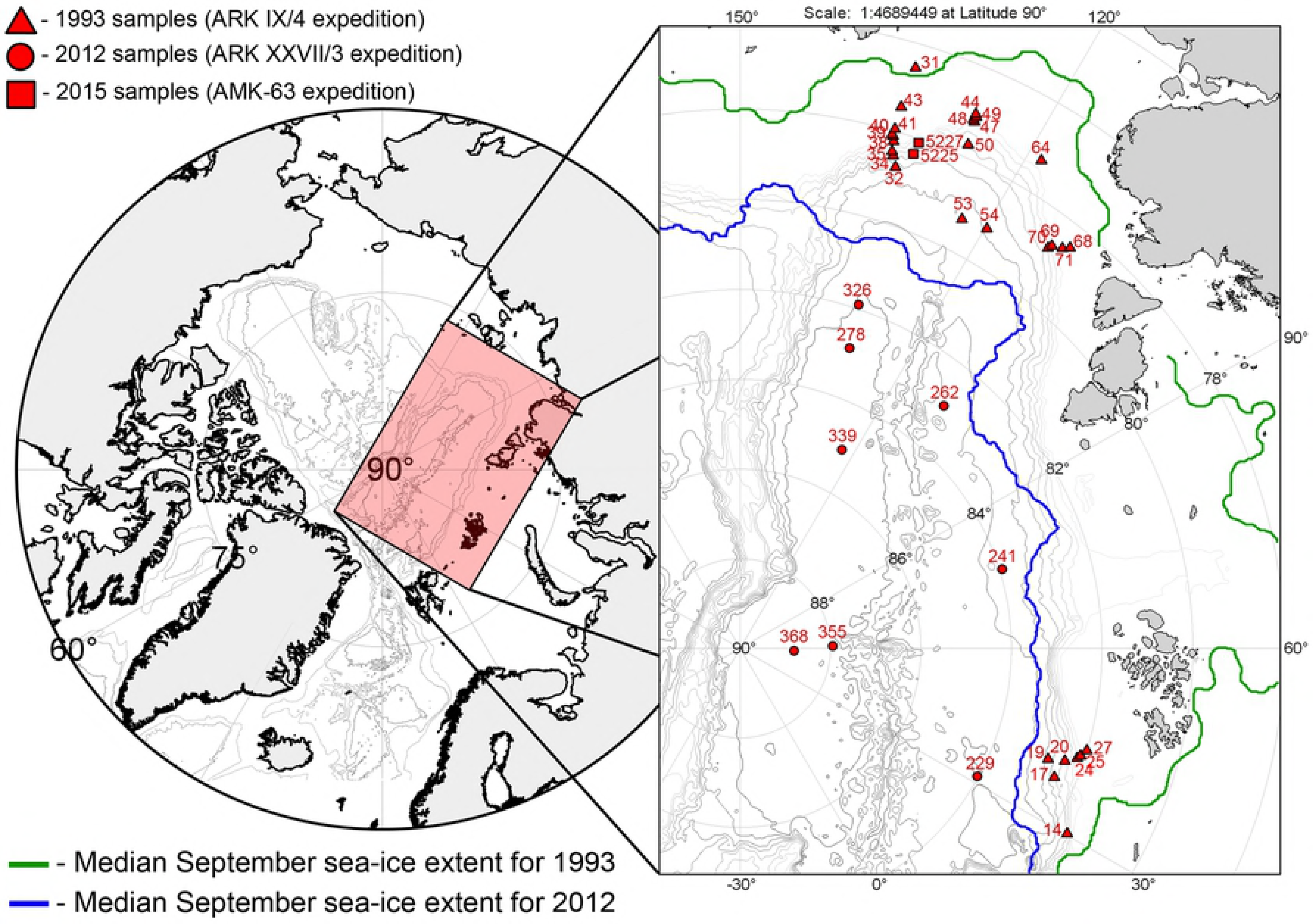
Sampling area, mean sea-ice extent and the stations.

Identification of macrofauna specimens to the lowest possible taxonomical level was made in the laboratory with the help of taxonomic experts (see ‘Acknowledgements’).

### Statistics

All specimens were counted and weighted (wet weight). Density and biomass were calculated per square meter. Dominant taxa in terms of abundance were distinguished for each station. The similarity between samples was estimated using the quantitative index of Bray-Curtis. We used the non-transformed data as a measure of taxonomic abundance because of the evenness across the most abundant taxa. Similarity matrixes were used for the cluster and multidimensional scaling (MDS) analyses. Clusters were generated using the UPGMA algorithm. Results were verified by ANOSIM analysis to reveal different station groups. Species richness was estimated as a total number of species at each station. Taxonomic diversity was estimated using the Shannon-Wiener index and the Hurlbert rarefaction index for 100 individuals. The species-individuals accumulation curves for the fauna were plotted. Spearman’s rank correlation was calculated between the environmental variables and community/species characteristics. A canonical correspondence analysis (CCA) was performed to estimate the contribution of the sediment parameters, depth and ice-extent to taxonomic composition [35]. The differences between different years of sampling were tested by the similarity percentages routine (SIMPER). For consistency in depth zonation, only the stations taken at depths <2000 m were tested for decadal variation.

Statistical analyses were performed using the Microsoft Excel 2007, Primer V6 [36] and PAST3 [37] software.

## Results

In total 10117 individuals (ind.) were obtained from 37 stations newly sorted for this study in a standardized manner. Most of thespecimens were sampled in 1993 (9443 ind. in the 38-3234 m depth range; Tab. 1). New samples added 101 ind. from the 2012 campaign (3432-4381 m depth range) and 573 ind. from the campaign in 2015 (2020-2390 m depth range). The specimens were attributed to 440 taxa. The density of macrobenthos varied from 12 ind. m^−2^ (St. 229) to 24957 ind. m^−2^ (St. 31). The biomass ranged from 0.06 g ww m^−2^ (St. 229) to 650.53 g ww m^−2^ (St. 25). The list of identified taxa with density and biomass values at each station is available in doi.pangaea.de/10.1594/PANGAEA.890152.

### Depth zonation of macrobenthos communities

The non-metrical cluster analysis and multidimensional scaling revealed five distinct groups of macrobenthic communities distinguished by depth Table 2, Fig 2). Between all zones, the maximum species overlap was 16% (Table 2). The community composition was significantly different (*p*-value of 0.001-0.022). The outer SHELF (38-54 m), UPPER SLOPE (73-1216 m), LOWER SLOPE (1991-3054) and PLAIN (3236-4381 m) groups were distinguished based on the Bray-Curtis similarity level (Fig 2a). We further distinguished as the fifth group the MID-SLOPE cluster from the UPPER SLOPE zone, because of the taxonomical peculiarity of the former (Fig 2, Table 3). The main dominants by density are shown in Table 3. Diversity characteristics of each group are given in Table 4. Species richness, contribution of each taxon to the total number of individuals, and to the total biomass within each station group are shown in Fig 3.

**Fig 2.**
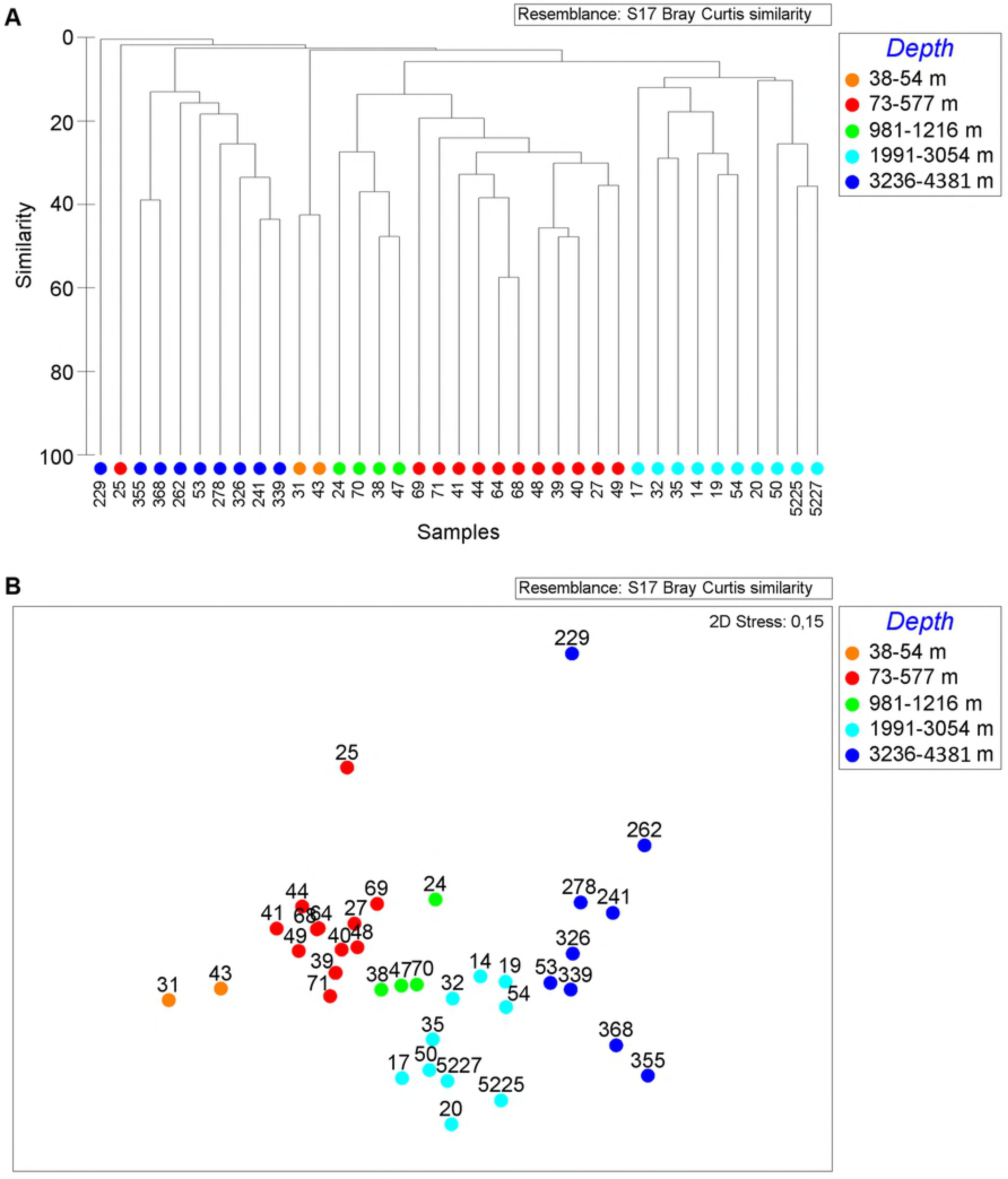
Cluster analysis (A) and non-metric multidimensional scaling plot (B) of all stations using the Bray-Curtis similarity index. Color indicates station group distinguished by depth. Orange - SHELF-group; red - UPPER SLOPE-group; green - MID-SLOPE-group; light-blue - LOWER SLOPE-group; dark-blue - PLAIN-group.

**Fig 3.**
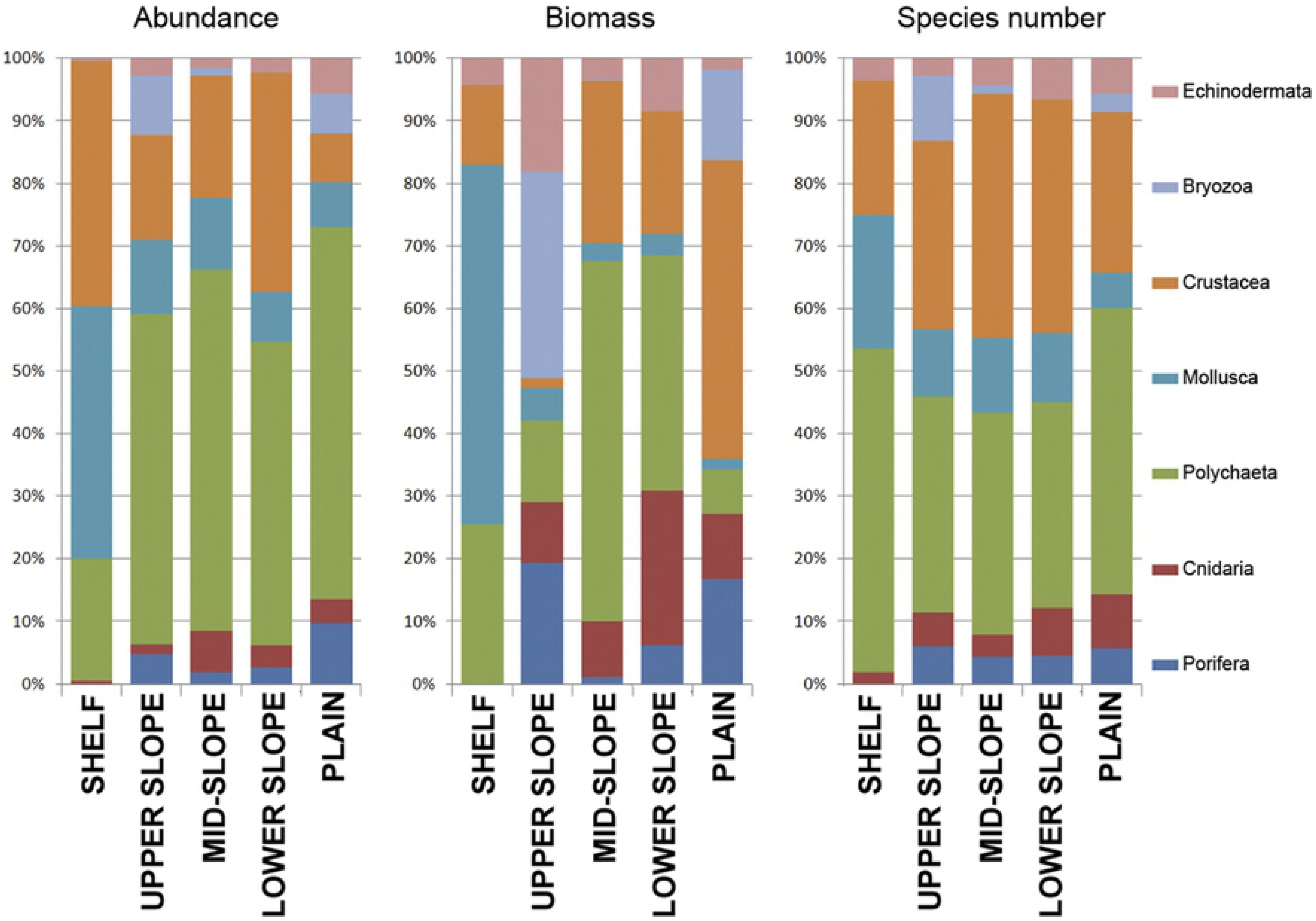
Percentage diagrams of total abundance, biomass and number of species per taxonomic group.

**Table 2.** Results of the ANOSIM analysis (abundance data). MID-SLOPE-group is included in the UPPER SLOPE-group.

| Groups pairs | R statistic | Significance level (%) | Possible permutations | Actual permutations | % Shared taxa |
| --- | --- | --- | --- | --- | --- |
| SHELF – UPPER SLOPE | 0.879 | 0.7 | 136 | 136 | 9.41 |
| SHELF – LOWER SLOPE | 1.000 | 1.5 | 66 | 66 | 3.68 |
| SHELF – PLAIN | 0.838 | 2.2 | 45 | 45 | 3.15 |
| UPPER SLOPE – LOWER SLOPE | 0.781 | 0.1 | 3268760 | 999 | 11.76 |
| UPPER SLOPE – PLAIN | 0.946 | 0.1 | 490314 | 999 | 4.55 |
| LOWER SLOPE – PLAIN | 0.700 | 0.1 | 43758 | 999 | 15.97 |

**Table 3.**
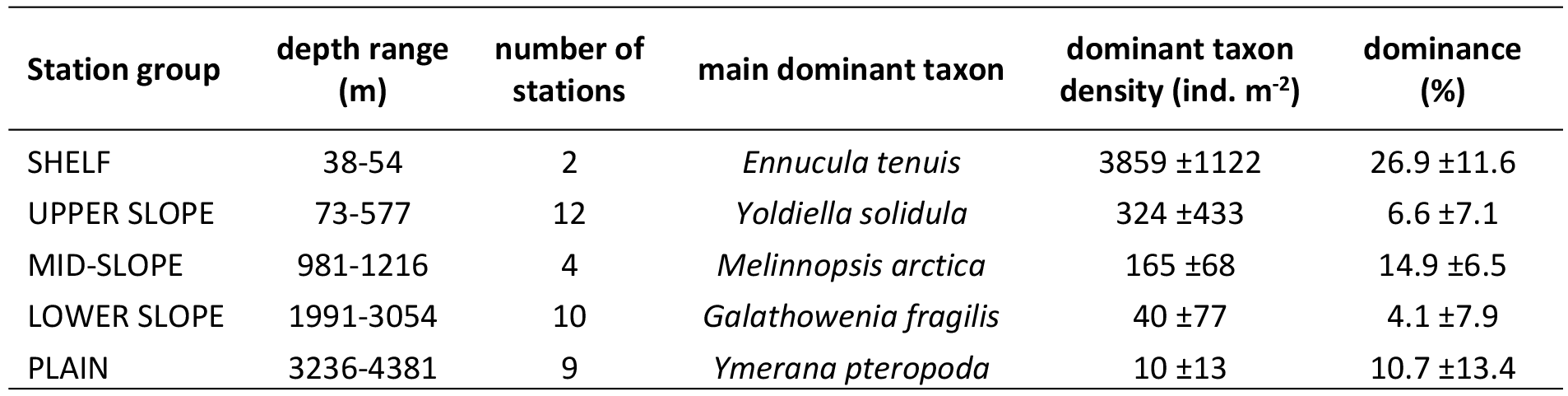
Dominant taxa of the station groups. Depth range, number of stations in each station group, density of the dominant taxa and the mean dominance level with standard deviation are shown.

| Station group | depth range (m) | number of stations | main dominant taxon | dominant taxon density (ind. m<sup>-2</sup>) | dominance (%) |
| --- | --- | --- | --- | --- | --- |
| SHELF | 38-54 | 2 | *Ennucula tenuis* | 3859 ±1122 | 26.9 ±11.6 |
| UPPER SLOPE | 73-577 | 12 | *Yoldiella solidula* | 324 ±433 | 6.6 ±7.1 |
| MID-SLOPE | 981-1216 | 4 | *Melinnopsis arctica* | 165 ±68 | 14.9 ±6.5 |
| LOWER SLOPE | 1991-3054 | 10 | *Galathowenia fragilis* | 40 ±77 | 4.1 ±7.9 |
| PLAIN | 3236-4381 | 9 | *Ymerana pteropoda* | 10 ±13 | 10.7 ±13.4 |

**Table 4.** Diversity characteristics for different depth ranges. Mean values of species number, abundance, biomass, Pielou evenness, Hurlbert rarefaction index for 100 individuals and Shannon-Wiener index with standard deviation are shown.

| Station group | Species per station | abundance (ind. m<sup>-2</sup>) | biomass (g ww m<sup>-2</sup>) | Pielou evenness | ES (100) | Shannon index |
| --- | --- | --- | --- | --- | --- | --- |
| SHELF | 35 ±23 | 16847 ± 11467 | 220.28 ±251.21 | 0.56 ±0.01 | 14.7 ±77.2 | 1.91 ±0.45 |
| UPPER SLOPE | 83 ±16 | 5293 ±2259 | 104.01 ±194.95 | 0.83 ±0.04 | 38.8 ±3.7 | 3.66 ±0.19 |
| MID-SLOPE | 64 ±32 | 1172 ±306 | 6.19 ±4.10 | 0.85 ±0.04 | 34.3 ±10.3 | 3.40 ±0.50 |
| LOWER SLOPE | 24 ±13 | 608 ±320 | 2.57 ±2.79 | 0.86 ±0.08 | 18.6 ±8.1 | 2.53 ±0.54 |
| PLAIN | 9 ±8 | 77 ±66 | 0.59 ±1.07 | 0.92 ±0.04 | 8.5 ±7.6 | 1.65 ±0.77 |

The SHELF group was characterized by the high dominance of bivalves *Portlandia arctica* and *Ennucula tenuis* and the ophiuroid *Ophiocten sericeum*. The biomass and density in this complex were the highest respectively, mostly owing to the bivalves (Fig 3, Table 4).

The UPPPER SLOPE group encompassed 12 stations. Dominant species in this group were the bivalve *Yoldiella solidula* and the polychaetes *Prionospio cirrifera* and Cirratulidae. Other notable species were the polychaetes *Proclea graffi* and *Notoroctus oculatus* and the cumacean *Ektonodiastylis nimia.* The biomass and density within this group were slightly lower than in the SHELF group, but diversity was the highest (Table 4). Polychaetes were the most abundant and diverse taxon, whereas the bryozoans and echinoderms (mainly the ophiuroids *Ophiacantha bidantata* and *Ophiocten sericeum*) contributed most to the biomass (Fig 3).

The MID-SLOPE group (four stations) was strongly dominated by the polychaete *Melinnopsis arctica* in terms of both the abundance and biomass. Other important species included the polychaete *Tharyx* sp., the bivalve *Yoldiella annenkovae*, the scyphozoan *Stephanoscyphus* and the sipunculid *Nephasoma diaphanes*. Crustacea was the most diverse taxon in this group (Fig 3). All diversity characteristics within the MID-SLOPE group, including the species number per station, density, biomass, ES (100) and Shannon index, were significantly lower than those in the UPPER SLOPE group (with the exception of Pielou evenness) (Table 4).

The LOWER SLOPE group included 10 stations. This group was rather uneven compared to others, and dominant species were different across stations, suggesting a substantial biodiversity variation at this water depth. The most abundant taxa included the polychaetes *Galathowenia fragilis*, *Ophelina opisthobranchiata* and *Terebellides* cf. *atlantis* and the tanaids *Pseudosphyrapus serratus* and *Pseudotanais* aff. *affinis*.

The PLAIN group (ten stations) was the least diverse among all station groups. The abundance, biomass, Shannon index and ES (100) were the lowest, whereas the Pielou evenness was the highest. Polychaeta was the most abundant and diverse taxon. Crustacea was more notable in terms of biomass, owing to the shrimp *Hymenodora glacialis* present in the catch at St. 53. This station taken near the Laptev Sea slope had the highest abundance, biomass and species number among the PLAIN group stations. The dominant taxa at stations of this group were different and included the polychaetes *Ymerana pteropoda* and *Anobothrus laubieri* and the sponge *Thenea abyssorum*. Other abundant taxon was the small bryozoan *Nolella* sp. Notable in samples of 2012 were scattered remains of semi-degraded ice-algae, apparently sedimented colonies of the diatom *Melosira arctica*.

Two stations, 25 and 229, were distinctly separate from other groups revealed by cluster analysis (Fig 2). Station 25 of the northern Barents Sea slope within the depth range of the UPPER SLOPE group (534 m) differed from other stations by the prevalence of suspension-feeders, including Porifera, Hydrozoa, Serpulidae, Bryozoa and the ophiuroid *Ophiacantha bidantata*. The multi-box corer sample at St. 229 taken in the Nansen Basin at the depth of 4008 m (the PLAIN group depth range) contained only one colony of the bryozoan *Nolella* sp. and an ovum of an unknown invertebrate.

In general the biomass and density of macrobenthos distinctly decreased with depth from the shallow-most stations to the deepest (Fig 4 a,b; Table 4). The results of the linear regression test for the abundance and biomass values are given in Table 5. The number of species, the Hurlbert rarefaction and the Shannon-Wiener index showed the maximum at the shelf break at depths ~100-300 m (Fig 5 a,b,c).

**Fig 4.**
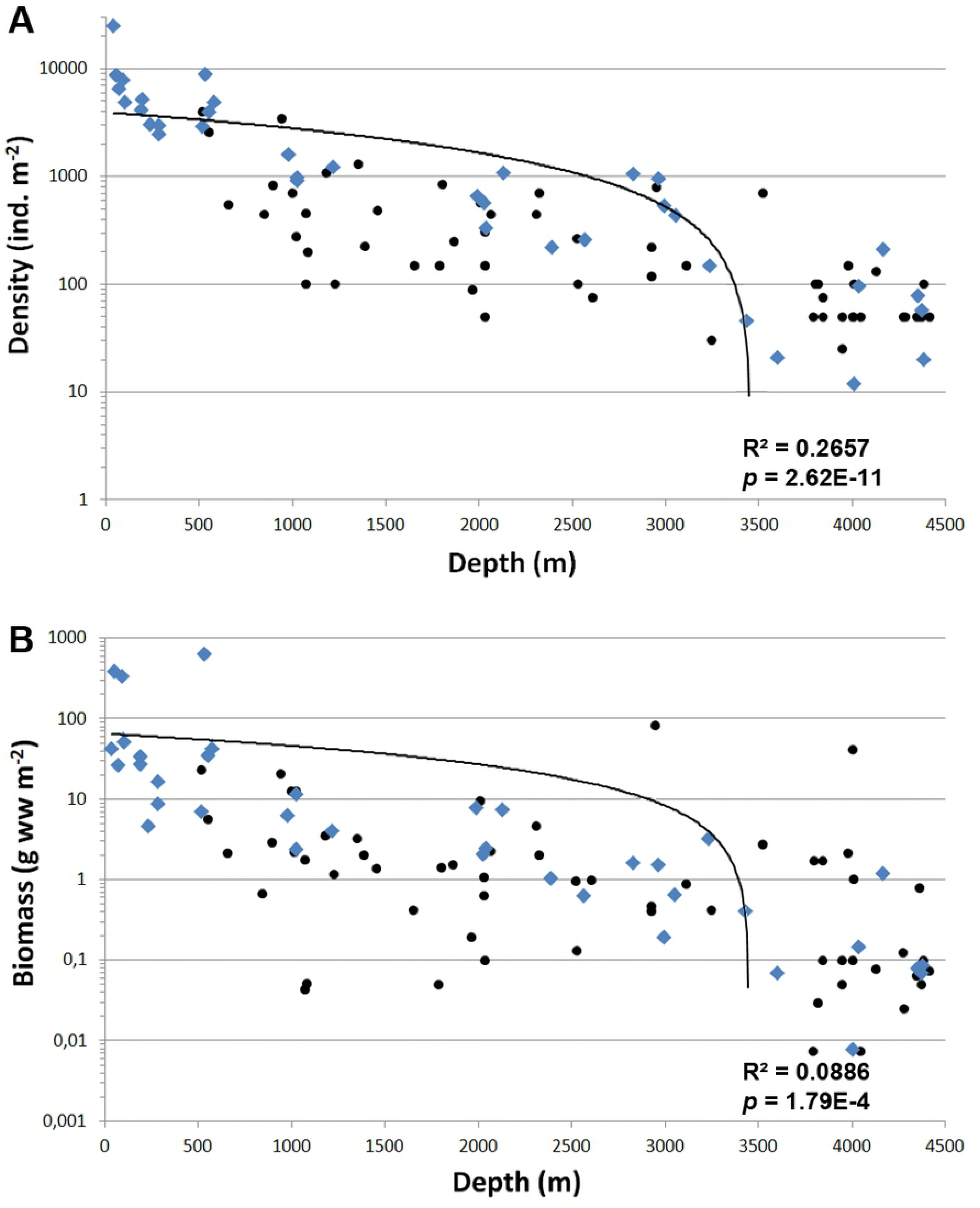
Abundance (A) and biomass (B) at stations in relation to depth. Blue rhombuses - our data; black dots - data from [6,28]. Lineal trend lines with R^2^ and *p* values calculated for all samples are added.

**Fig 5.**
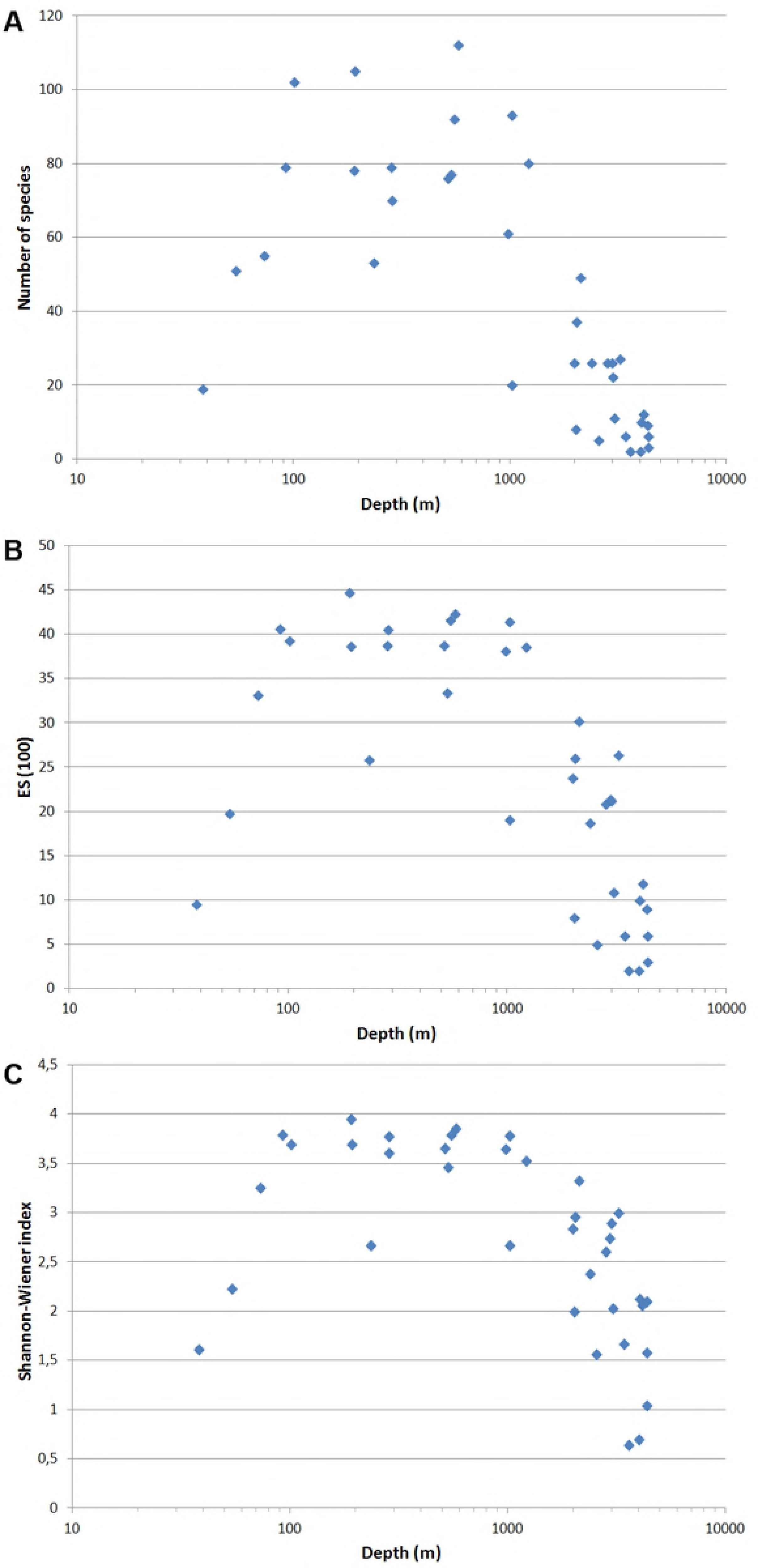
Number of species (A), Hurlbert rarefaction (B) and Shannon-Wiener index (C) at each station in relation to depth (logarithmic scale).

**Table 5.** Results of the linear regression test for the abundance and biomass values related to depth. Data from 1991 [28], 1993, 1997 [6] and 2012-2015 are shown.

|  |  | Number of samples | R<sup>2</sup> | p-value |
| --- | --- | --- | --- | --- |
| Abundance | all samples | 96 | 0.2657 | 2.62E-11 |
|  | 90-th samplings | 86 | 0.2601 | 1.93E-10 |
|  | 2000-th samplings | 10 | 0.5315 | 0.0047 |
| Biomass | all samples | 96 | 0.0886 | 1.79E-04 |
|  | 90-th samplings | 86 | 0.0852 | 3.55E-04 |
|  | 2000-th samplings | 10 | 0.6079 | 0.0032 |

### Environmental factors governing macrobenthos distribution

Environmental data available for the sample collections of 1993 and 2012 allowed to test the correlation of total abundance of macrofauna with total chloroplastic pigments, indicating phytodetritus deposition, and with bacterial abundance in the surface sediments (Table 6). Of the individual taxa, the abundances of the tanaid *Pseudotanais affinis*, scaphopod *Siphonodentalium laubieri* and polychaete *Ophelina opisthobranchiata* correlated significantly in abundance with chloroplastic pigments, bacterial abundance and organic carbon concentration (Table 6). The results of canonical correspondence analysis among the deep stations (>2000 m, LOWER SLOPE and PLAIN groups) are shown in Fig 6. Only those stations were chosen for which environmental variables were available for both years, 1993 and 2012. Stations 19, 20, 35 and 50 of the depth range 1991-2993 m demonstrated higher values of phaeopigments, bacterial abundance and organic carbon, and higher density of tanaid *Pseudosphyrapus serratus*, amphipod *Neohela monstrosa* and polychaete *Terebellides* cf. *atlantis*. The deep 2012 samples and the deepest 1993 station (53) showed low chloroplast pigments values, but higher densities of polychaetes *Ymerana pteropoda, Anobothrus laubieri* and bryozoan *Nolella* sp. (Fig 6). There was no direct correlation of these samples with the ice coverage, community composition and the abundance/biomass of distinct taxa.

**Table 6.** Spearman’s rank correlation between the community/species characteristics and the environmental/sediment parameters. Correlated pairs with p<0.05 are shown.

| Correlated variables | | Spearman's correlation | $p$ (uncorr.) |
| --- | --- | --- | --- |
| Community characteristics/species | Environmental/sediment parameter |  |  |
| Abundance total | Depth | -0.80 | 0.0004 |
| Abundance total | Php | 0.78 | 0.0005 |
| Abundance total | CPE | 0.72 | 0.0026 |
| Abundance total | Bact ab | 0.88 | 1.84E-05 |
| Biomass total | Depth | -0.74 | 0.0015 |
| *Pseudotanaïs affinis* biomass | Depth | -0.81 | 0.0003 |
| *Pseudotanaïs affinis* biomass | Php | 0.78 | 0.0006 |
| *Pseudotanaïs affinis* biomass | CPE | 0.70 | 0.0036 |
| *Pseudotanaïs affinis* biomass | Bact ab | 0.81 | 0.0003 |
| *Siphonodentalium laubieri* biomass | TOC | 0.70 | 0.0037 |
| *Ophelina opisthobranchiata* biomass | TOC | 0.72 | 0.0025 |

For environmental/sediment parameters see doi.pangaea.de/10.1594/PANGAEA.890152.

**Fig 6.**
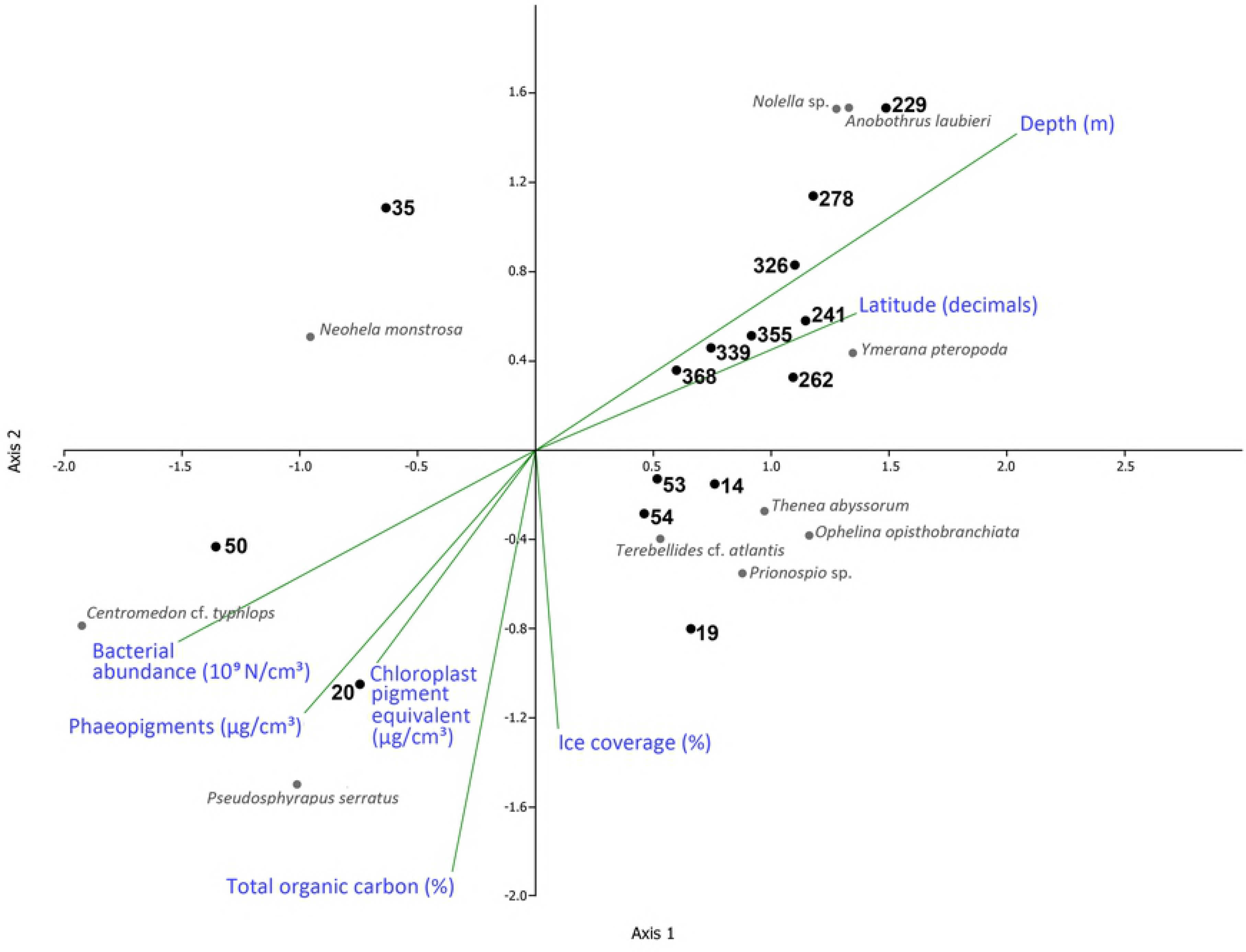
Results of canonical correspondence analysis (CCA) of the LOWER SLOPE and PLAIN station groups. Ten most abundant taxa are shown. Axis 1 significance level - 0.281; axis 2 significance level - 0.231.

Species-individual accumulation curves for each station group are shown in Fig 7. The curves tend to reach the saturation in the SHELF and UPPER-LOWER SLOPE groups, whereas in the PLAIN group increasing the number of samples could significantly increase the total species numbers (Fig 7).

**Fig 7.**
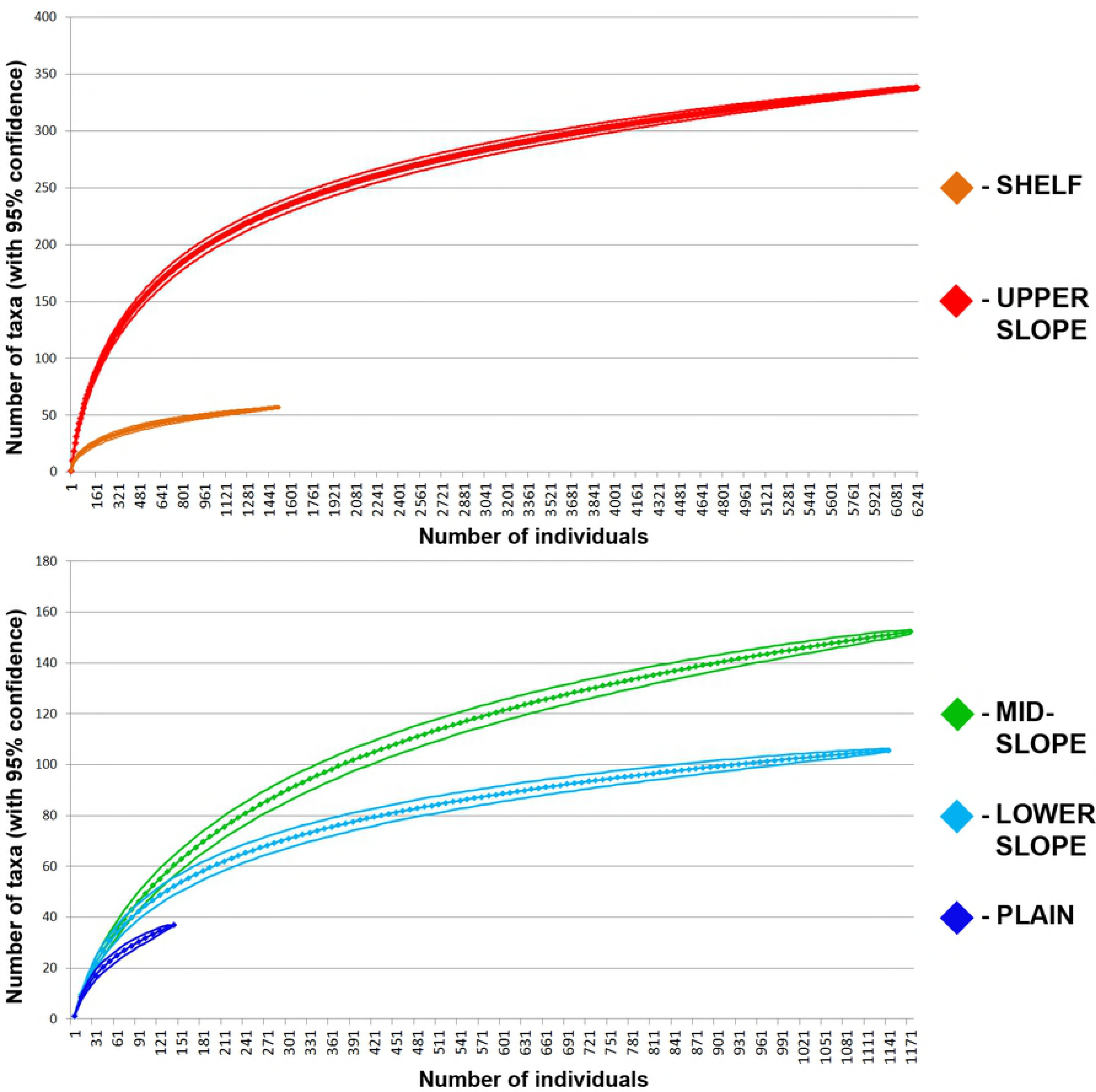
Species-individual accumulation curves with 95% confidence calculated separately for SHELF, UPPER SLOPE, MID-SLOPE, LOWER SLOPE and PLAIN station groups.

### Temporal trends

To assess temporal trends, we selected 19 stations from all three expeditions with some regional overlap (depth >2000 m). The mean abundance of these taxa from 1993 and 2015 samples with the standard deviation are shown in Fig 8. Some differences in the abundance of certain taxa were revealed by SIMPER-analysis, but the standard deviation overlapped, so that no reliable conclusions about temporal changes could be made.

**Fig 8.**
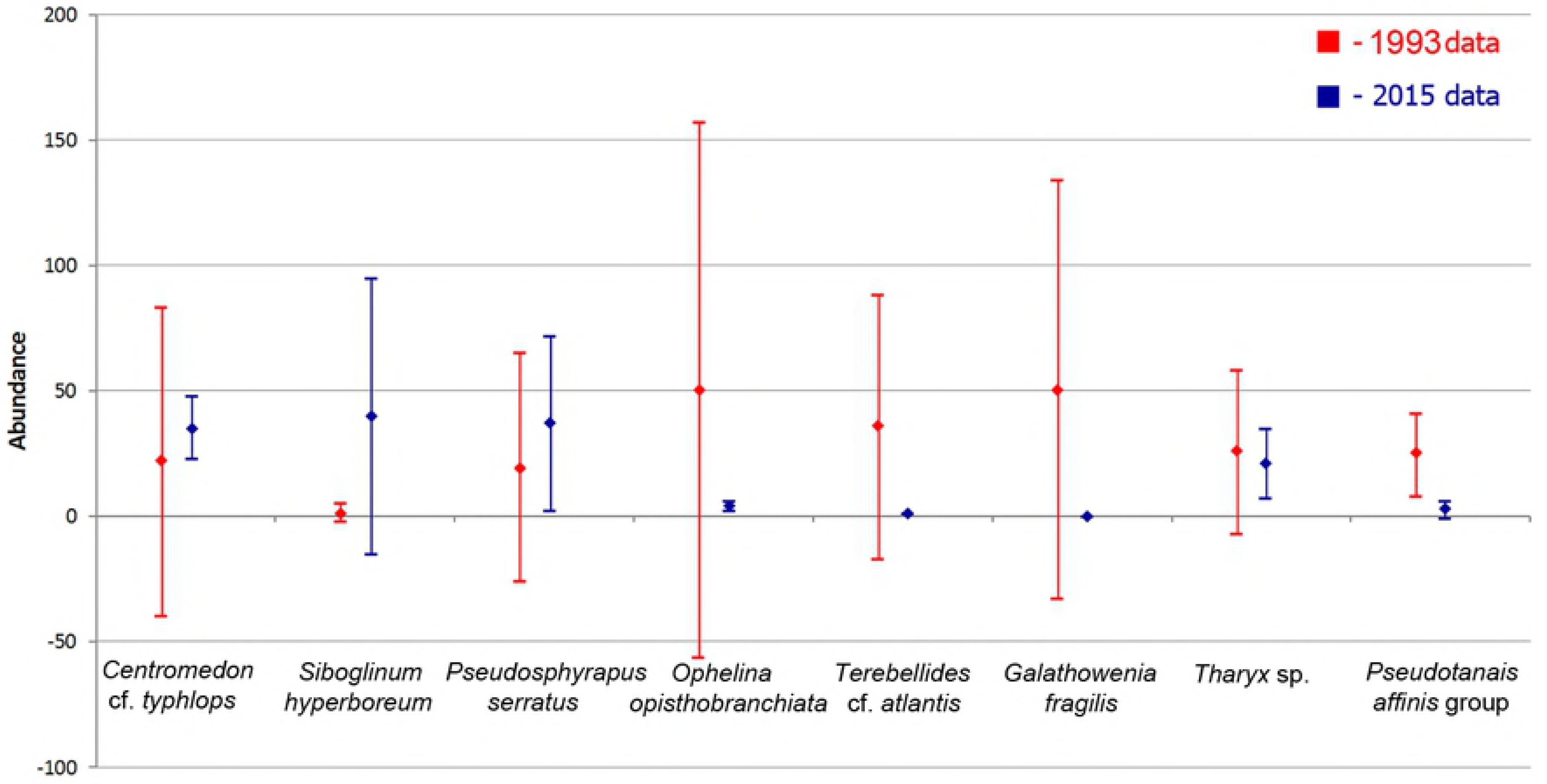
Abundance (mean values with standard deviation) of the species revealed by the SIMPER-analysis of 1993 and 2015 samples taken deeper than 2000 m.

When limiting to total biomass and abundance and including the data of [5] and [6] for the depth zone 2000-4500 and comparing all data, no significant differences between the samples within these two decades (1991-1997 vs 2012-2015) could be detected (Fig 9).

**Fig 9.**
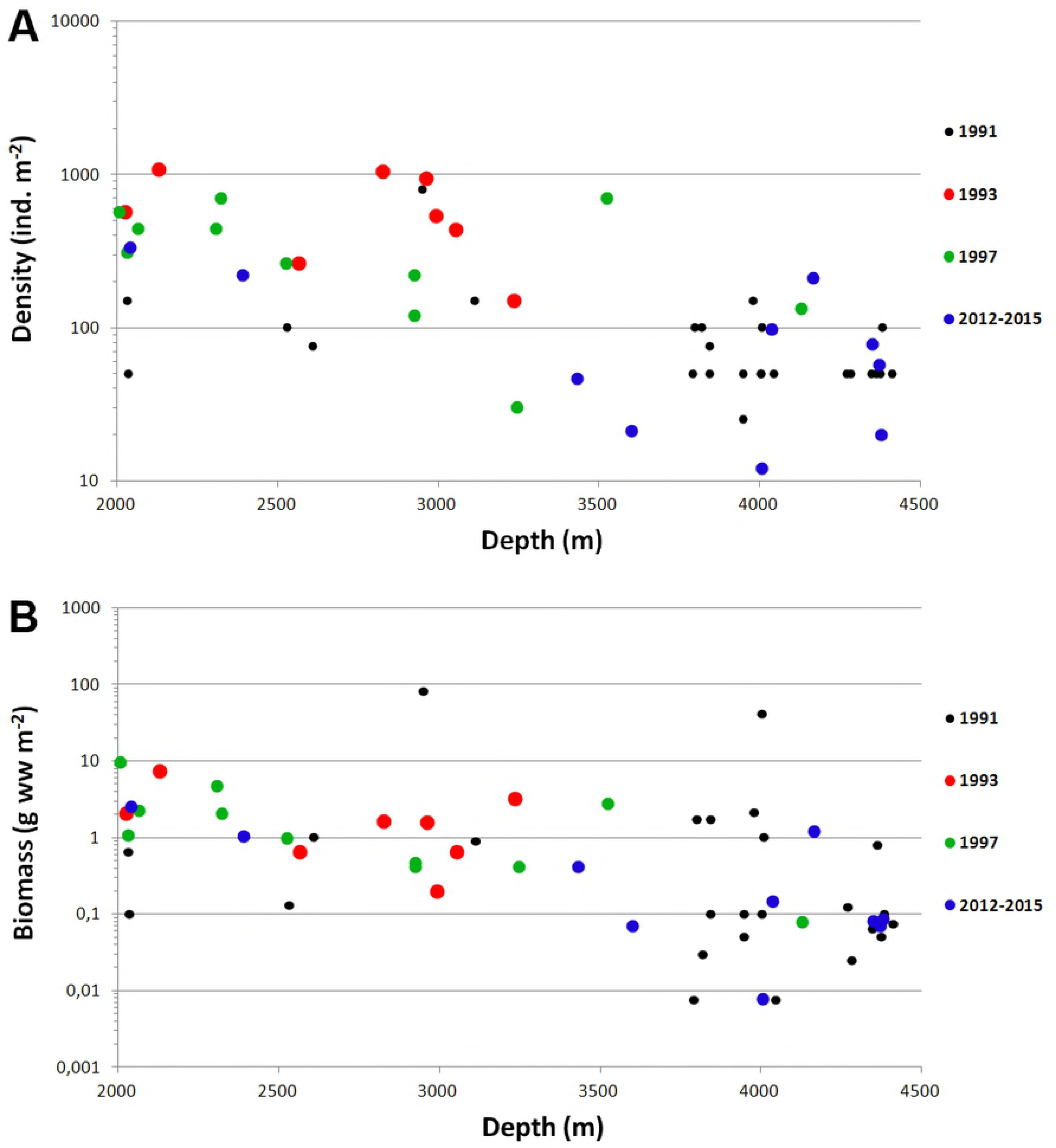
Abundance (A) and biomass (B) at stations in relation to depth (from 2000 to 4500 m). Data from 1991 [28], 1993, 1997 [6] and 2012-2015 are shown.

## Discussion

### Macrobenthos of the Central Arctic Ocean is controlled by food availability

Similar to results of previous investigations that addressed factors influencing macrobenthos abundance and composition in other oceans, the present study revealed the substantial decrease of density and biomass between the seasonally ice-free shelves and the year round ice-covered basins [5,27,28]. We summarized published data on the biomass and abundance for the depth ranges used in our study (Table 7). It can be concluded that the ice-covered Central Arctic shows some of the lowest macrobenthos abundances and biomasses for the Northern Seas and the North Atlantic.

**Table 7.** Abundance and biomass of macrofauna at different geographical sites and different depth ranges. The values are arranged by latitude.

| Region | Depth Ranges (m) |  |  |  |  | Reference |
| --- | --- | --- | --- | --- | --- | --- |
|  | 40-70 m | 70-600 m | 600-1300 | 1300-3000 | 3000-4200 |  |
|  | Abundance (ind./m²) |  |  |  |  |  |
| This study | 16847 ± 11467 | 5293 ±2259 | 1172 ±306 | 608 ±320 | 77 ±66 | This study |
| Central Arctic | - | - | 256 ±148 | 200 ±271 | 58 ±49 | 13 |
| Central Arctic | - | 2495 ±356 | 706 ±95 | 1042 ±854 | 349 ±341 | 70* |
| Fram Strait, Central Arctic | - | 4048 | 1294 ±1516 | 430 ±266 | 281 ±346 | 6* |
| Lomonosov Ridge | - | - | 58 ±8 | 22 ±26 | 6 ±4 | 11 |
| Canada Basin | - | - | 50 ±26 | 78 ±65 | - | 3* |
| Canada Basin | - | 4300 | 1867 ±383 | 1156 ±1061 | 113 ±62 | 71 |
| Kara Sea slope | - | 1988 ±1242 | 755 | 271 ±205 | - | 16 |
| Laptev Sea slope | 1220 ±732 | 2195 ±2924 | 420 ±150 | 516 ±434 | 252 ±133 | 11 |
| Franz-Josef Land | 6635 ±657 | 2785 ±1408 | - | - | - | 39 |
| Off Spitsbergen, 79°N | - | 2902 ±2180 | - | 646 ±240 | - | 38 |
| Off Mauritania, 18-21° N | - | - | - | 3984-5432 | 204-1840 | 40* |
| Goban Spur, 49° N | - | 4785-7980 | 2882-6169 | 1413-1655 | 542-648 | 41* |
| Biomass (g ww/m²) |  |  |  |  |  |  |
| This study | 220.3 ±251.2 | 104.0 ±194.9 | 6.19 ±4.10 | 2.57 ±2.79 | 0.59 ±1.07 | This study |
| Central Arctic | - | - | 1.80 ±0.46 | 0.90 ±1.11 | 0.38 ±0.53 | 13 |
| Central Arctic | - | - | 0.55 ±0.39 | 0.40 ±0.19 | 0.14 ±0.20 | 70* |
| Fram Strait, Central Arctic | - | 23,7 | 12.44 ±7.09 | 2.89 ±3.01 | 0.89 ±1.25 | 6* |
| Lomonosov Ridge | - | - | 0.29 ±0.27 | 0.31 ±0.61 | 0.013 ±0.013 | 11 |
| Canada Basin | - | - | 0.11 ±0.07 | 0.64 ±1.08 | - | 3* |
| Kara Sea slope | - | 643.8 ±995.8 | 28,1 | 2.35 ±0.05 | - | 16 |
| Laptev Sea slope | 82.3 ±51.2 | 33.4 ±34.2 | 3.68 ±2.97 | 1.75 ±1.35 | 0.34 ±0.47 | 15 |
| Franz-Josef Land | 3465 ±827.5 | 404.6 ±462.5 | - | - | - | 39 |
| Off Spitsbergen, 79°N | - | 38.7 ±14.5 | - | 2.20 ±1.50 | - | 38 |
| Off Mauritania, 18-21° N | - | - | - | 2.37 ±1.19 | 0.05 ±0.01 | 40 |
| Goban Spur, 49° N | - | 33.3-71.2 | 1.60-51.20 | 2.30-4.00 | 2.40-4.20 | 41* |

\*- the values include meiobenthos

In the shelf and slope areas similar values of abundance and biomass were reported for Laptev Sea [11,15], Fram Strait [38], Nansen Basin and Yermak Plateau [6]. Lower values were shown for the Lomonosov Ridge [11]. For the Arctic Ocean higher values of biomass were shown for the Kara Sea upper slope [16] and Franz-Josef Land [39] mostly due to sponges and other suspension-feeders. In the Atlantic Ocean much higher values of abundance were shown for the Mauritania slope [40] and for the Goban Spur area [41].

The global bathymetrical trends of biomass in the ocean were analyzed by [42]. The authors reported a biomass decrease from ~4 mg C m^−2^ on the Arctic shelf to <2.5 mg C m^−2^ in the Arctic deep sea, with a higher rate of decrease in the Arctic compared to the temperate and tropical areas of Atlantic and Pacific Oceans. The present study adds new data for the Central Arctic deep basins, confirming this trend of a more rapid decrease, most likely explained by low productivity and export flux.

The biomass and density values we observed for the abyssal stations (0.008-1.212 g ww m^−2^) agree with previously obtained data from other parts of the deep-sea Arctic [3,6,12]. [13,14] reported higher biomass and density values from the western parts of the Nansen and Amundsen Basins. However, this may result from samples contaminated by boreal shallow-water organisms, such as the polychaete *Magelona* sp., the amphipod *Jassa marmorata*, the bivalve *Spisula elliptica* etc. [13,14]. [15] suggested that these organisms could inhabit the pipeline system of a research vessel. After excluding those shallow-water species from the analysis, the biomass and density values appeared similar to those in our results (Table 7).

The overall decrease in abundance and biomass is connected with the food availability - the most important factor in determining the structure and function of benthic communities [43]. The amount of chloroplast pigments is commonly used as indicator of the particle flux and food availability in benthic ecosystems. It was shown that the amount of pigments decreases from the shelf edge towards abyssal depths in the Arctic Ocean [6,43]. In our study the abundance was strongly correlated with the phaeopigments and CPE values (Table 6).

Among 10 stations taken from >3200 m depth, the highest biomass and density were registered at the southern-most St. 53 taken closest to the continental slope base. All other stations of this depth group were situated on the abyssal plain at some distance from the slope. Benthic biomass and abundance declined in the direction from the continental slope base towards poorer central abyssal areas. This effect in the Central Arctic can be explained by increasing ice-cover, limiting primary productivity [6,33,43].

During the *Polarstern* ARK27-3 cruise numerous deposits of fresh diatom algae *Melosira arctica* were observed on the seafloor including areas of all 2012 stations except for St. 229 [17]. This diatom was observed forming meter-long filaments on the under-ice surface in remarkable densities. Calculations showed that in the extreme year 2012 marked by the largest sea-ice melt recorded to date, algal deposition had reached an average 9 g C per m^2^ with *Melosira* contribution >85% of the carbon export. It was suggested that this significant increase in algal production could have resulted from recent climate changes affecting sea-ice and snow cover thickness, an important limiting factor for primary production [6,17,44].

These food falls of *Melosira* were shown to attract some of opportunistic megafaunal organisms, first of all holothurians and ophiuroids [17]. Degraded parts of *Melosira* were discovered in some of 2012 samples. However, no evidence of direct macrobenthic response to mass algal input to the seafloor was observed yet, and polychaete abundances were very low in 2012 as previously. However, the overall significantly positive correlation between the total abundance of macrobenthos and phaeopigments volume in our samples suggests that an overall increase in the sedimentation of phytodetritus is likely to change faunal abundance and composition in the future. Key indicator species for an increase in algal input could be tanaid *Pseudotanais affinis*; its biomass is strongly correlated with phaeopigment concentrations (Table 6, Fig 8).

### Community similarity is high across all depth zones

Previous studies of macrofauna community composition suggest a high proportion of endemic species in the Central Arctic with a large overlap in species between shallow and deep regions due to uniform temperature regime and other biogeographical factors [5,45]. Here we further investigated the hypothesis that despite large variations in abundance, a significant proportion of communities would be distributed independent of water depth in the Arctic. We used species richness and species diversity (the number of species per number of individuals) for comparison.

Species distribution in the Arctic is partially explained by the “polar emergence” of fauna (review in [46]). [46] distinguish two different phenomena termed in the literature the “polar emergence”: the extension of bathymetric ranges of recent bathyal and abyssal species in the Arctic to the shelf depths (i.e. the rise of deep-water species to shallow depths) (I) and the “evolutionary emergence and submergence” related to the colonization of different bathymetric zones in the history of shelf fauna (II). Both phenomena may increase the number of species at lower shelf - upper slope depth in the Arctic Ocean. Thus, [46] demonstrated that among 133 species occurring deeper 2000 m in the Arctic, 27% can be found at depths between 200 and 1000 m and 41% at depths from 0 to 200 m (the most shallow depth of occurrence). This effect may also increase the number of species at upper slope - lower shelf depths. Our data show that 121 species found in LOWER SLOPE and PLAIN zones (e.g. deeper 2000 m, see doi.pangaea.de/10.1594/PANGAEA.890152) contribute 45% to the community composition of MID-SLOPE zone, 47% to UPPER SLOPE and 5% to SHELF. However, beyond the similarity in presence/absence, the macrofauna composition showed a substantial dissimilarity of the different depth zones.

In the present study the macrobenthos clustered significantly according to water depth, despite some differences in region, time and washing protocols (Fig 2). Exceptions were one station of a high current regime at the shelf edge, and one almost devoid of fauna (Nansen Basin). The statistical test with ANOSIM supported that differences between the depth ranges were significant, and higher than differences in species composition between particular samples from different areas (i.e. Barents and Laptev seas slope). The only other extensive study of benthic community composition in the Central Arctic also detected substantial water depth zonation [11]. This study was based on the samples of the RV *Polarstern* expedition ARK-XI/1 in 1995 and the IB *Oden* expedition Arctic Ocean-96 in 1996. Seven station groups were revealed by [11] on the eastern slope of the Laptev Sea, the Lomonosov Ridge and adjacent deep-sea parts of the Amundsen and Makarov Basins. The SHELF group distinguished in our study corresponds with the SHELF group shown by [11]. This group is characterized by benthic communities dominated by the bivalves *Ennucula tenuis* and *Portlandia arctica*, which are widely distributed at shallow depths of the Laptev and East-Siberian Seas [47,48]. The diverse UPPER SLOPE group in our study agrees with the HANG (i.e. Slope) and RAND (i.e. Edge or Margin) station groups of [11]. This group is characterized by the *Ophiocten sericeum* and *Ophiopleura borealis* communities typical for the lower shelf - upper bathyal depths of the eastern Arctic [49–51]. In our study only two stations were taken on the outer shelf at depths <70 m (stations 31 and 43 forming the SHELF group) with a total sampled area 0.115 m^2^. The UPPER SLOPE group corresponding to the lower shelf - upper slope depths (73-577 m) consists of twelve stations with the total area sampled equivalent to 1.357 m^2^. It is obvious that increasing the number of stations from these depths would make conclusions more reliable.

The MID-SLOPE group in our study matches the *Melinnopsis arctica* community described by [1] from the bathyal depths of the northern Kara Sea and the Lomonosov Ridge. [11] included stations from the similar depth range (981-1216 m) in the RAND group. The diverse LOWER SLOPE group in our study possibly corresponds to the RAND and RUCKEN (i.e. Ridge) communities described by [11].

The PLAIN group in our study matches the TIEF (i.e. Deep) group described by [11]. The species-individual accumulation curve for the PLAIN group suggests that the number of species will sharply increase with additional samples (Fig 7). At the same time, since the density of benthos in the deep-sea Central Arctic is lower than in other parts of the ocean [6,28], it can be assumed that the small sampling area of generally used quantitative gears (0.20-0.25 m^−2^) is not enough for the accurate estimation of regional species diversity.

Several taxa found in the PLAIN group have not been recorded in the Central deep-sea Arctic so far. The polychaete *Ymerana pteropoda* was previously known from one locality north from Svalbard [52], one locality in the Canada Basin north from Alaska [53] and two localities north-east off Iceland [54]. In our samples this species was present and even dominant in terms of density at five stations. Thus the known distribution of this species was extended significantly. In the genus *Nolella* (Bryozoa) the abyssal species were known only from the North Atlantic [55,56]. These peculiar bryozoans of a tentatively new species were present in two of our samples. These minute organisms could have been overlooked during previous investigations owing to their small size (1-3 mm) and dense silt coverage of zooids masking them among sediment particles.

We further investigated trends in macrobenthos diversity with water depth in comparison to other ocean areas. For the North Atlantic it was shown that the peak of species richness and diversity occurs at depths 2000-3000 m [26,57,58]. In the Goban Spur area of the north-east Atlantic species diversity increased with water depth from 208 m (ES(100) = 29) to 4115 m (ES(100) = 68) [25]. At the same site, the species richness per sample showed a different pattern, since the number of species is dependent on sample area, numerical density and species diversity [25]. [38] found that in Fram Strait, both species diversity and richness decreased with depth (depth range from 203 to 2977 m, ES(100) declined from 43 to 14). In the present study we detected a parabolic pattern of species diversity and richness change with depth, with the species maximum occurring at 100-300 m (ES(100) = 39 ±4), i.e. shallower than the bathymetric range investigated by [38] in Fram Strait.

A number of published studies on macrofauna on the Siberian shelf showed an increase of species richness and diversity from the upper shelf to the shelf edge [48,59–62]. Besides the explanation of this peak in diversity and richness by the “polar emergence” theory explained above, we also suggest that food availability by higher productivity of the ice margin zone is a key factor. In the years 1980 - 2008 the seasonal ice margin was mostly found across the zones UPPER SLOPE and MID-SLOPE. Only recently it retreated as far as beyond the BASE of the slope into the PLAIN [63]. It remains an important task to continue surveys of potential benthic community shifts in response to the retreating ice-margin in the Eurasian sector of the Arctic Ocean.

The comparably lower macrobenthos richness and biomass on the Siberian Arctic shelf could be a regional phenomenon, related to the enormous river run-off, potentially impacting ocean stratification and thereby food availability. The Ob, Yenisei and Lena rivers contribute almost 1700 km^3^ of fresh water per year into the Kara and Laptev Seas, which is about 60% of the total Arctic rivers annual discharge [64,65]. For the Kara Sea it was shown that brackish waters generated by the rivers Ob and Yenisei add to the stratification of the shelf sea and form a thick and extensive layer on the surface. The layer spreads in the form of giant lenses and can be traced even along the east coast of the Novaya Zemlya Islands [66]. The layer of brackish waters reduces nutrient transport in summer [67]. This may explain the lower species richness of benthos on the shelf compared to the shelf edge. In the Laptev Sea a peak in primary production related clearly to the border of the freshwater lens was found near the shelf-edge above the depths 100-200 m [68]. Another possible reason of the lower diversity on the shallower parts of Siberian shelf could be the sea-ice scraping. Large furrows on the sediment surface left by sea ice can be traced down to 15-20 m depth [69]. However, the influence of ice scraping on benthic macrofauna has not been studied in our study area.

### Temporal changes

In the present study we synthesized and tested temporal and regional differences in benthic data in terms of Panarctic perspective ([5,28], this study). However, it remains currently impossible to assess potential temporal trends of the macrobenthos in response to the rapid sea ice retreat of this decade, due to the low number of benthic samples available from the Arctic Ocean (Fig 8). Comparison of the total abundance and biomass data revealed no differences between different years of samplings (Fig 9). However, the set of stations taken in the Central Arctic deeper than 2000 m is rather small. Repeated surveys from the same areas are needed for the correct analysis of the potential changes in macrobenthos community composition in the Nansen and Amundsen basins. Furthermore, a taxonomical revision and standardizing of previously published species lists [11,15,16,70] by one pool of taxonomical experts is required.

## Conclusions

The present work has demonstrated that on the slope of the Barents and Laptev Seas and the abyssal of the Nansen and Amundsen Basins, the macrobenthos communities group according to water depth. Five station groups were distinguished based on species composition analysis in the depth range from 38 m to 4381 m: 1) group on the shelf (38-54 m); 2) group on the lower shelf and upper slope; 3) group on the mid-slope (981-1216 m); 4) group on the lower slope (1991-3054 m); and 5) group on the abyssal plain (3236-4381 m). These groups partly correspond to the groups described by [11]. The distribution of the groups is strongly correlated with depth and group boundaries do not overlap bathymetrically. The polychaete *Ymerana pteropoda* and the bryozoan *Nolella* sp. were recorded in open areas of Nansen and Amundsen Basins for the first time. Rarefaction analyses predict that the existing diversity of slope and basin macrobenthos has not been adequately sampled yet.

Decrease of density and biomass of macrobenthos with depth was confirmed for the Central Arctic Basin. The depth-related parabolic pattern of species diversity with the maximum at 100-300 m was shown for the Arctic for the first time. Temporal changes caused by thinning and retreating sea ice, which could potentially enhance primary productivity and export fluxes, are impossible to identify given the sparse sampling of the Arctic Ocean floor.

## Acknowledgements

Thanks are due to the captains, crews and shipboard parties of the RV *Polarstern* expeditions in 1993 (ARK IX/4) and 2012 (ARK XXVII/3) and the RV *Akademik Mstislav Keldysh* expedition in 2015 (AMK-63) for multiple help with the work on board and obtaining samples. The authors thank Dr. Eike Rachor for collecting the 1993 samples, Dr. Renate Degen for the 2012 samples and Dr. Sergey Galkin for the 2015 samples. Our special thanks to taxonomic experts Dr. Dorte Janussen, Dr. Nataliya Budaeva, Dr. Vyacheslav Labay and Dr. Alexandr Mironov for help in identifying sponges, polychaetes, amphipods and echinoderms respectively. We acknowledge Dr. Christina Bienhold for providing access to the environmental datasets. We also thank Dr. Bodil Bluhm, Dr. Vadim Mokievsky and Dr. Ingrid Kröncke for informative discussions and for the data they gave us.

